# A novel method for multi-scale recording of intracranial EEG through dynamic alteration of electrode surface area

**DOI:** 10.1101/2022.01.05.475164

**Authors:** Kavyakantha Remakanthakurup Sindhu, Duy Ngo, Hernando Ombao, Joffre E. Olaya, Daniel W. Shrey, Beth A. Lopour

**Affiliations:** Department of Biomedical Engineering, University of California, Irvine, Irvine, CA; Department of Statistics, Western Michigan University, Kalamazoo, MI; Statistics Program, King Abdullah University of Science and Technology, Saudi Arabia; Division of Neurosurgery, Children’s Hospital of Orange County, Orange, CA; Department of Neurosurgery, University of California, Irvine, Irvine, CA; Division of Neurology, Children’s Hospital of Orange County, Orange, CA; Department of Pediatrics, University of California, Irvine, Irvine, CA

## Abstract

Intracranial EEG (iEEG) plays a critical role in the treatment of neurological diseases, such as epilepsy and Parkinson’s disease, as well as the development of neural prostheses and brain computer interfaces. While electrode geometries vary widely across these applications, the impact of electrode size on iEEG features and morphology is not well understood. Some insight has been gained from computer simulation studies and experiments in which signals are recorded using electrodes of different sizes concurrently in different brain regions. Here, we introduce a novel method to record from electrodes of different sizes in the exact same location by changing the size of iEEG electrodes after implantation in the brain. We first present a theoretical model and an *in vitro* validation of the method. We then report the results of an *in vivo* implementation in three human subjects with refractory epilepsy. We recorded iEEG data from three different electrode sizes and compared the amplitudes, power spectra, inter-channel correlations, and signal-to-noise ratio (SNR) of interictal epileptiform discharges, i.e., epileptic spikes. We found that iEEG amplitude and power decreased as electrode size increased, while inter-channel correlation increased with electrode size. The SNR of epileptic spikes was generally highest in the smallest electrodes, but 39% of spikes had maximal SNR in medium or large electrodes. This likely depends on the precise location and spatial spread of each spike. Overall, this new method enables multi-scale measurements of electrical activity in the human brain that facilitate our understanding of neurophysiology, treatment of neurological disease, and development of novel technologies.

## 1. Introduction

Intracranial electroencephalography (iEEG) is an invasive technique that measures electrical activity of the brain. It is used for the diagnosis, monitoring, and treatment of neurological diseases, such as epilepsy (1) and Parkinson’s disease (2). It has also been critical to the development of devices such as neural prostheses (3) and brain computer interfaces (BCIs)(4). iEEG measurement is done using subdural grids or strips of electrodes placed on the surface of the cerebral cortex or depth electrodes inserted into brain tissue. The iEEG electrodes are sensitive to the electric signal in their immediate vicinity, which is produced by the aggregate post-synaptic activity of cortical neurons (5). The voltage measured by an electrode is thought to reflect the average potential distribution under its uninsulated contact area (6–8). The signals recorded by a subdural grid depend on a number of factors, including the impedance, geometry, and spacing of the electrodes (5).

Because the number of neurons whose electrical activity contribute to the iEEG signal is proportional to the electrode contact area, electrode size is an important factor in measurements. A multitude of electrodes of different geometries are used for intracranial EEG measurement. Microwires with diameters as low as 12 μm have been used for *in vivo* single unit recordings (9). Micro-electrocorticogram (ECoG) electrodes, which have potential uses in both BCI and clinical applications, have diameters in the range of 10 μm to several hundreds of micrometers (10). Standard clinical macroelectrodes have exposed diameters that range from 0.86 mm to 3 mm (11). Therefore, it is critical to understand the precise relationship between electrode size and iEEG measurement to better interpret and compare the results of studies with different methodologies.

*In silico* studies of the effect of electrode size on iEEG signal characteristics present conflicting pictures. Nelson and Pouget (12) developed a physical model that predicted electrodes with larger surface area would have higher correlation between them. Their model also suggested that, as the voltage profile underneath the electrode becomes more inhomogeneous (as it would with increasing surface area), electrodes with different contact areas are more likely to measure different average values. Ollikainen, Vauhkonen (6) simulated rectangular electrodes with surface areas that varied from 1.5 cm^2^ to 5 cm^2^, measuring electrical potential from a single source. They showed that smaller electrodes had more sensitivity to localized voltage differences than larger electrodes. This is consistent with the idea that each electrode measures the average potential of the underlying tissue; therefore, using larger electrodes results in loss of spatial information. Furthermore, these simulations showed that the current distribution on the surface of the electrode is non-uniform and concentrated at the boundaries, with the distribution becoming more complex when larger electrodes are used. Contrary to this, the model developed by Suihko, Eskola (13) suggested that changing the electrode size will cause only small changes in the sensitivity distribution and is therefore not a key factor in iEEG measurements.

A number of studies have analyzed neuronal action potentials and the effect of electrode size on their measurement (14–17). Anderson et al. posit that, in the context of action potentials, as the size of the electrode increases, the “listening sphere” increases, but the signal-to-noise ratio (SNR) decreases (15). However, a study by Ward et al. in an animal model found no significant difference in the action potential SNRs for implanted micro-electrode arrays (MEAs) of different surface areas (16). The results of such studies can be confounded by differences in the electrode coating and other techniques to lower the electrode impedance, independent of the electrode diameter.

In contrast, there are few *in vivo* studies analyzing the effect of electrode size on general iEEG characteristics, such as amplitude or waveform morphology (18, 19). Some studies in humans have reported that smaller electrodes are advantageous, but consensus on this issue has not been reached. In applications like BCI and neuro-prosthetic devices, the ability to accurately classify neural signals associated with different cognitive tasks is of utmost importance. Studies have shown that smaller electrode size and higher grid density enable the recording of signals from smaller spatial scales, making them more suitable for these applications (19–21). In human studies of epilepsy, various electrode sizes have been used to measure high frequency oscillations (HFOs), a candidate biomarker for epileptogenic brain tissue. HFOs are highly localized, transient iEEG events characterized by high-amplitude 80-500 Hz oscillations. Using human intracranial EEG, Chatillon et al (22) analyzed HFOs recorded with electrodes of different sizes and found that the difference in recordings was not clinically relevant. On the other hand, Worrell et al. reported that smaller electrodes recorded more HFOs than larger electrodes, particularly in the 250-500 Hz frequency range (23). Another study by Boran et al. found that HFO measurement is aided by the use of a high-density electrode grid with a smaller surface area than a standard subdural grid (24). Knowledge of the relationship between electrode size and iEEG biomarker features would inform epilepsy surgery and invasive monitoring, with the potential to improve patient outcomes.

Therefore, the goal of this study was to directly measure the impact of electrode surface area on iEEG signal characteristics in the human brain. Previous work in humans has relied on simultaneous iEEG recordings using electrodes of different sizes, with each electrode implanted in a different location. In those cases, it is not clear if the resulting differences are due to electrode size or regional differences in brain activity. An alternative methodology is to record sequential iEEG measurements using electrodes of multiple sizes, placed over the exact same region of neural tissue. However, because implantation of intracranial electrodes is an invasive procedure with inherent risk to the patient, this presents logistical and ethical challenges. Here, we present a solution to this problem: a method to alter the size of an intracranial recording electrode after implantation. This enables multiscale measurements from a single region of the brain, allowing for a more direct comparison of iEEG signals recorded using electrodes of different sizes. We also examine how electrode size affects basic iEEG properties, such as power spectrum and amplitude, and we explore the effect on the morphology of interictal epileptiform discharges (also called epileptic “spikes”), which is a common electrographic event in epilepsy patients.

## 2. Theoretical basis for altering electrode size via electrical shorting

Our method to alter the surface area of an implanted grid electrode involves electrically connecting adjacent electrodes together (via physical shorting) to generate a range of effective surface areas. For each individual electrode, the recorded signal reflects the average voltage across the uninsulated surface; therefore, connecting two adjacent electrodes will report the average of the two individual electrodes, equivalent to doubling the electrode surface area (7, 8). In this way, we can alter electrode size while recording data from the exact same region of neural tissue within the framework of standard clinical care.

To provide a theoretical basis for this approach, we used a widely accepted electrical circuit model of the metal electrode (Figure 1A) (25, 26). In Figure 1B, two cortical surface electrodes (with impedances Z_e1_, Z_e2_) are each connected to an amplifier (with impedances Z_a1_, Z_a2_). The electrodes sense independent voltage sources in the neural tissue (V_s1_, V_s2_), which interact through the impedance of the brain tissue (Z_b1_, Z_b2_) and a shunt impedance between the two brain regions (Z_12_).

**Figure 1:**
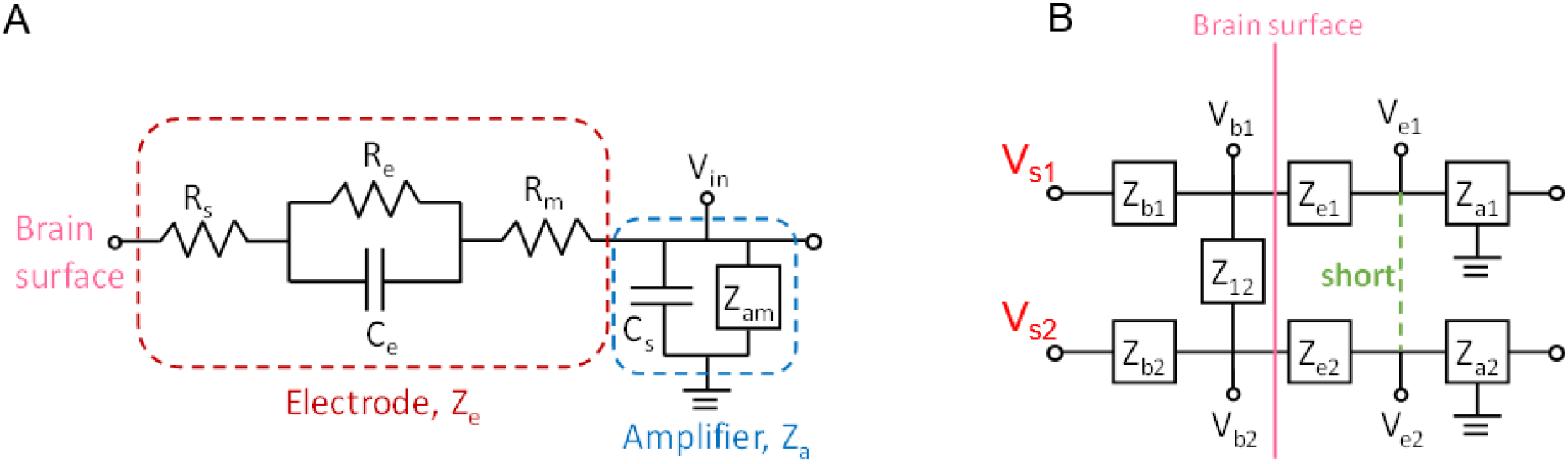
(A) Circuit model of a single electrode (red box) connected to an ideal amplifier (blue box). (B) Circuit model of two electrodes (Ze1, Ze2) on the surface of the brain that can be shorted together (green dashed line) to simulate an electrode with twice the surface area.

In the case with no shorting, each electrode measures the voltage in the underlying tissue, ***V***_*e1*_ ≈ ***V***_*b1*_ and ***V***_*e*2_ ≈ ***V***_*b*2_, assuming the input impedance of the amplifier is large. When the two electrodes are shorted, they measure a common voltage ***V***_*e*_ given by the equation:

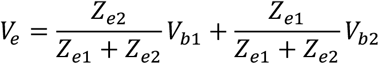

The full derivation of this equation can be found in the Appendix. If we assume the electrode impedances *Z*_*e1*_ and *Z*_*e2*_ to be equal, we find that the voltage measured at the surface is

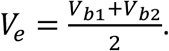

This result suggests that, when two electrodes are shorted together, the voltage measured by the amplifiers is a linear combination of the voltages sensed by each individual electrode (V_b1_ and V_b2_). Because signals measured by iEEG electrodes are thought to reflect the average neural activity underneath them, this averaged activity should be equivalent to that sensed by a larger electrode covering the same cortical area as the two smaller electrodes. Therefore, physically shorting two adjacent electrodes can increase the effective surface area, thus providing a means to investigate the same region of neural tissue with electrodes of different sizes.

## 3. Methods

### 3.1 *In vitro* validation experiment

We performed an *in vitro* experiment to test the prediction from the circuit model that the signal recorded when electrodes are shorted together is equal to the average of the individual electrode signals (Section 2). A disc of agar gel was used as the substrate because it has been shown to mimic both the structural and electrical properties of brain tissue (27). The gel was mixed with water and NaCl to achieve a conductivity of ~0.5 S/m, to approximately match that of brain tissue (28). A standard 4-by-4 grid of electrodes was placed in the center of the disc (Figure 2A). Bipolar electrical stimulation was applied to the gel substrate using a 1-by-6 electrode strip, and a second 1-by-6 strip was placed on the opposite side for use as an electrical reference. The stimulus was a sine wave of 80Hz frequency and 350µV amplitude. The sinusoidally oscillating dipole created an electric field in the substrate that was sensed by the 4-by-4 grid of electrodes (Figure 2B). The signals from those electrodes (n=16, “single” electrodes) were recorded using Biopac amplifiers and were sampled at 2kHz.

**Figure 2:**
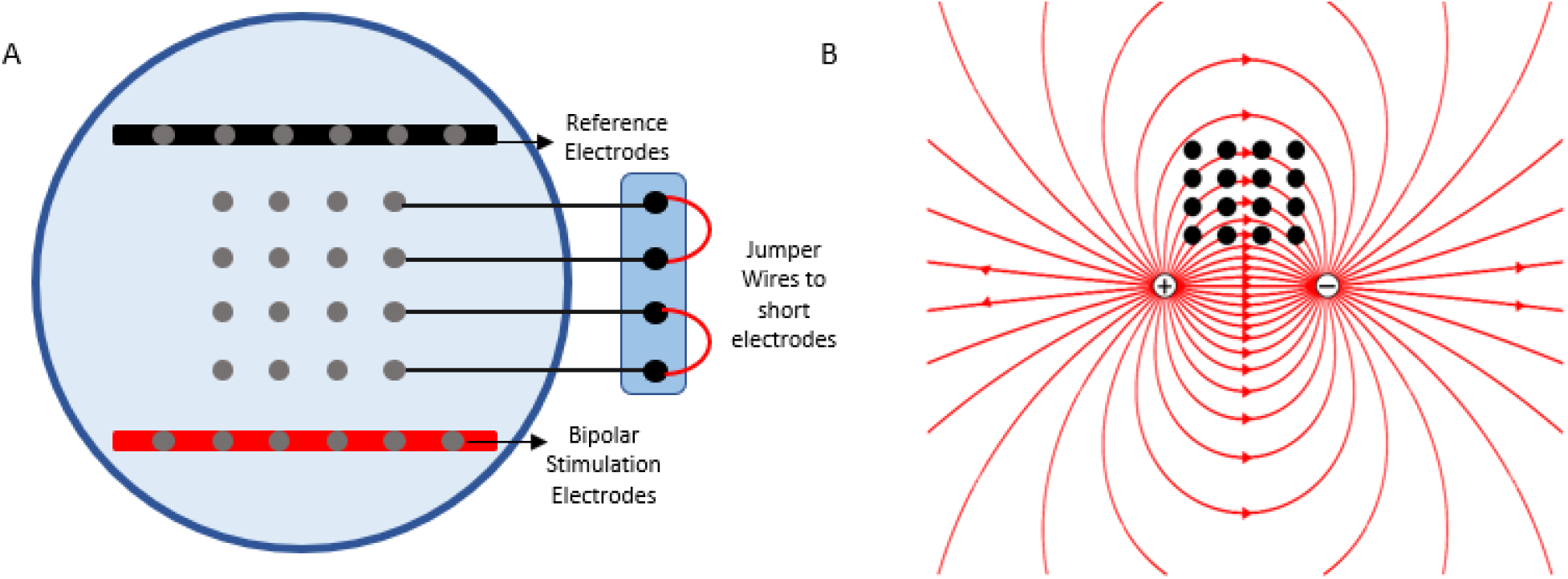
(A) Schematic of the agar gel disc with the 4×4 grid of electrodes, the 1×6 strip containing the reference electrode, and the 1×6 strip used for stimulation. An example is shown in which two pairs of electrodes are shorted at the amplifier using jumper wires. (B) The red lines represent the electric field lines of the dipole being sensed by the grid.

Then pairs of adjacent electrodes were shorted together using jumper wires connected at the junction with the amplifier (Figure 2A). The same bipolar stimulation was applied, and the resulting electric field was measured at each shorted electrode pair (n=8, “pair” electrodes). Finally, we repeated the stimulation and recording when 2-by-2 sets of four adjacent electrodes were shorted together (n=4, “quad” electrodes).

We compared the signals recorded by the single, pair, and quad electrodes. To validate the model, we also compared them to the signals obtained via mathematical averaging of the single electrodes, e.g., we averaged pairs of signals from the single electrode recording and compared those to the physically paired electrode signals. If the model is valid, mathematical averaging and physical shorting should produce the same signal. We used a rank-based non-parametric approach (Wilcoxon rank sum test) to test for the differences in correlation and power spectrum of signals obtained from the “pair” and “quad” configurations.

### 3.2 *In vivo* experiment

#### 3.2.1 Human data collection

The *in vivo* portion of this study was approved by the Institutional Review Board of the Children’s Hospital of Orange County (CHOC). Informed consent was obtained prior to involvement in the study. Three human subjects with medically intractable epilepsy were each implanted with a high-density 8×8 subdural grid of intracerebral EEG electrodes (Ad-Tech FG64C-MP03X-000) as part of phase 2 pre-surgical invasive monitoring. Each electrode had an exposed surface area of 1.08mm^2^ (we will refer to these as the “small” electrodes) and electrode spacing was 3mm center-to-center. The effective surface area was changed by electrically shorting adjacent electrodes in groups of two and four, thereby mimicking larger surface areas of 2.16mm^2^ (“medium” electrodes) and 4.32mm^2^ (“large” electrodes), respectively (Figure 3A). This was done by connecting jumper wires to the paired electrodes at the jack box outside the patient’s body (Figure 3B). The jack box combines the individual electrode wires into an integrated cable before connecting to the amplifier. A quick-release connector enabled rapid reconfiguration of the electrode shorting, minimizing disruption to the patient’s recording. The jack box and jumper wires were placed in a Faraday cage to minimize electrical interference.

**Figure 3:**
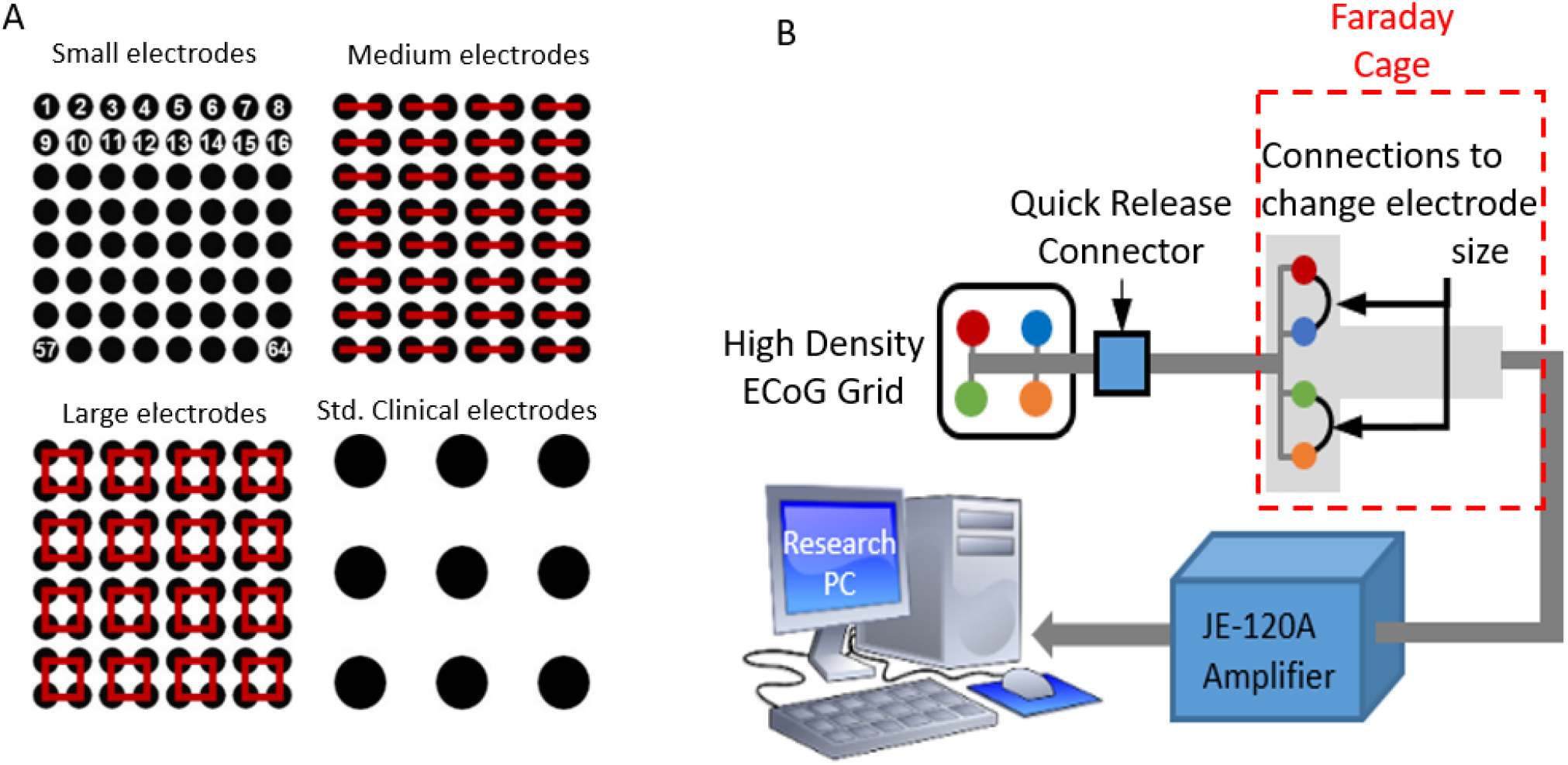
(A) Schematic that shows the shorting technique to create medium and large surface areas, as well as a scale drawing of a standard grid size, for comparison. (B) Experimental setup, showing an example in which a small 2×2 grid (red, blue, green, and orange electrodes) are shorted together in pairs (red-blue and green-orange) using jumper wires.

20-minute iEEG recordings were thus obtained for each of three different electrode surface areas (small, medium, large) from a single grid in a static brain location while the subjects were sleeping. The sampling rate was 5kHz, and the data were referenced to the common average of the 8×8 grid. The iEEG data were high pass filtered using a zero phase FIR filter at 1Hz and notch filtered at 60Hz, 120Hz, and 180Hz to remove electrical line noise before analysis.

Similar to the *in vitro* study, we compared the results to the theoretical circuit model by generating “simulated medium” electrode signals (consisting of the mathematical average of adjacent pairs of small electrodes) and “simulated large” electrodes (the mathematical average of four adjacent small electrodes). The averaging of signals was done on the raw iEEG data and the data were then re-referenced and filtered as described above. We compared the medium and large electrode recordings to their associated simulated signals.

#### 3.2.2 Data analysis

##### Correlation

For each subject, five one-minute segments of data spaced at least two minutes apart were used for this analysis. The data were band pass filtered into three frequency bands using a zero phase FIR filter: low frequency (1-30Hz), gamma-1 (30-60Hz), and gamma-2 (60-100Hz). In each frequency band, correlation values were calculated in five-second windows for every pair of channels that were at a center-to-center distance of 6mm for all three electrode sizes (n=96 for small, n=48 for medium, n=24 for large electrodes). Then the correlation values were averaged over all time windows for a given frequency band, electrode size, and electrode pair. For each subject, the samples of size-specific mean correlation values were compared across electrode sizes using a Wilcoxon rank sum test. To assess the baseline distribution of correlation values (under the null hypothesis of zero correlation), correlation between channels was calculated using surrogate data. The surrogate data *x*_*s*_(*t*) was obtained by applying a random circular time shift to the original data *x*(*t*) as follows:

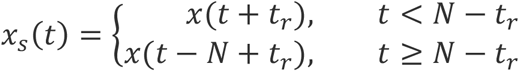

Where *t*_*r*_ is a random time point ranging from 1 to N-1.

##### Amplitude

The iEEG data from each of the three electrode configurations was bandpass filtered in the frequency range of 1-100Hz. The bandpass filtered EEG was then divided into 100 segments of five seconds each. For each 5-second segment, the root-mean-square (RMS) value of the amplitude was calculated. These samples of RMS amplitude values were compared across electrode sizes using a Wilcoxon rank sum test.

##### Power Spectrum

For every subject, power spectral density from 1-100 Hz was estimated for each of the 100 epochs of five seconds, for each electrode configuration. The following statistical analysis was done independently for each subject. To enable paired comparisons of the power spectra, the signals were grouped based on the large electrode configuration. For example, small electrodes 1, 2, 9, and 10 were compared to two medium electrodes (1 shorted to 2, and 9 shorted to 10) and one large electrode (1, 2, 9, and 10 shorted together) (see Figure 3). For each set of four small electrodes, the power spectra were estimated by using the Fourier periodogram which is the data-analogue of the spectrum defined on the fundamental frequencies. Since periodograms are quite noisy, they need to be smoothed in order to obtain a mean-squared consistent estimator (29). In some applications, it is more convenient to use log periodograms (rather than periodograms) because their variance is approximately constant across frequencies. Here, log periodograms were calculated and smoothed across frequencies using a moving average filter with a span of 5 data points (0.5 Hz). For each set of two medium electrodes, the two log periodograms were averaged and smoothed using a span of 10 data points (1 Hz). For the large electrodes, the log periodograms were used without averaging and were smoothed using a span of 20 data points (2Hz).

Thus, for each configuration, a set of 1600 log periodograms was obtained (16 signals x 100 epochs). To explore structures, patterns, and features in the sample of periodograms’ curves, we followed Ngo et al. (2015) and constructed a functional box plot (FBP) (31), a generalization of the classical pointwise boxplot. For each curve, a modified band depth (MBD) value is computed (32). This indicates whether or not a curve is covered by many pairs of curves in the data. Based on the ranks of MBD values, the FBP provides descriptive statistics, such as the functional median curve which has the highest MBD value.

##### Depth-based Permutation Test for the Power Spectrum

Let *F, G* and *L* be the distribution of log periodogram populations from three different settings (small, medium and large electrodes) with *n*_1_ = *n*_2_ = *n*_3_ = 1600. We propose a depth-based permutation test for our null hypothesis that the three populations of curves come from the same distribution, i.e., there is no difference in the distributions of the small, medium, and large electrodes. Let 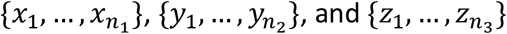, and 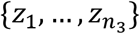 denote the three samples’ curves from distributions *F, G* and *L*, respectively.

Suppose that 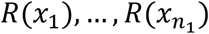 are the corresponding ranks of 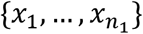, measured by comparison to the combined three samples of size *n*_1_ + *n*_2_ + *n*_3_. The test statistic is defined as 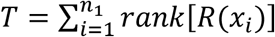, which is the sum of the MBD ranks in distribution *F*. Under the null hypothesis, *T* is the sum of *n*_1_ numbers that are evenly distributed between 1 and *n*_1_ + *n*_2_ + *n*_3_. If the alternative hypothesis is true, the sample *x*_*i*_ will be more outlying than the other samples, which implies that the depth values will be smaller, with correspondingly smaller ranks. Thus, a small value of *T* provides strong evidence to reject the null hypothesis. Since it is a challenging task to obtain the distribution of *T* under the null hypothesis in a case of three samples, we then carry out a permutation test to compute the p-values, which is as follows:

1. Permute electrode configuration labels (small, medium and large) among the samples in the combined set 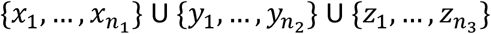 and denote the resulting samples of the *j*th permutation to be 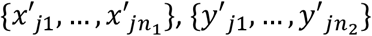, and 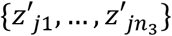 for *j* = 1, …, *J*.
2. For each permutation, we compute the test statistic 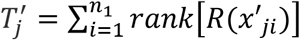
3. The p-value is approximated by 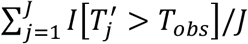 where *T*_*obs*_ is the observed value of *T* based on the original combined samples 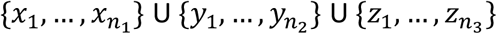, and *I* is the indicator function.

Because the different electrode configurations were recorded at different times, we also tested whether the power spectra were stable over time. For each configuration, we compared the power spectra in the first five minutes of the recording to those in the last five minutes, using five equally spaced 25-second intervals for each case. Depth based permutation testing was done as described above on the two sets of five curves in each scheme.

#### 3.2.3 Analysis of Interictal Spikes

We also wanted to characterize the impact of electrode size on the morphology of transient electrographic events. Because the study subjects had refractory epilepsy, we focused on interictal epileptiform discharges, i.e., interictal spikes. For this analysis, 20-minute segments of data were used, each one clipped from the long-term recording while the patient was sleeping, between midnight and 12:30am. Interictal spikes were manually marked in the iEEG data in the small electrode configuration for each subject under the supervision of a board-certified epilepsy specialist (DS). We then simulated each spike in the medium and large electrode configurations by averaging the corresponding small electrode data. We defined the SNR of each spike as the signal to background amplitude ratio. The amplitude of the spike was measured as the difference between the minimum and maximum voltages recorded over the duration of the spike. To calculate the background amplitude, a one-second interval around the spike, not containing the spike, was considered. The signal in this window was rectified, and the average of the rectified signal was defined as the baseline amplitude. The SNRs were compared across the three electrode configurations and each spike was classified into one of three types: type S, in which a small electrode had the highest SNR, type M, in which a medium electrode had the highest SNR, and type L, in which a large electrode had the highest SNR. The spatial spread, defined as the combined area of the electrodes in which the SNR of the signal exceeded 1.5 during the time of the spike, was also calculated for each spike.

## 4. Results

### 4.1 *In Vitro* Study: Physical shorting of electrodes is consistent with mathematical averaging, as suggested by the circuit model

The *in vitro* experimental results are shown in Figure 4. For the single, pair and quad electrodes, the measured signal maintained the original sinusoidal shape and 80 Hz frequency of stimulation (Figure 4A). The RMS amplitudes varied according to the electric field created by the bipolar stimulation (Figure 4B). For the single configuration, the highest amplitude was observed at the boundary of the grid, and the amplitude decreased for the inner electrodes. Amplitude also decreased as the vertical distance from the stimulating electrodes increased. Both results are consistent with the electric field lines in Figure 2B. When varying the physical shorting, we found that the amplitudes were highest for the smallest electrodes and decreased with an increase in electrode size. The RMS values from the mathematically averaged signals (Figure 4B, lower right) were not significantly different from the RMS values obtained by physically shorting the electrodes (Wilcoxon rank sum test, p-value>0.5).

**Figure 4:**
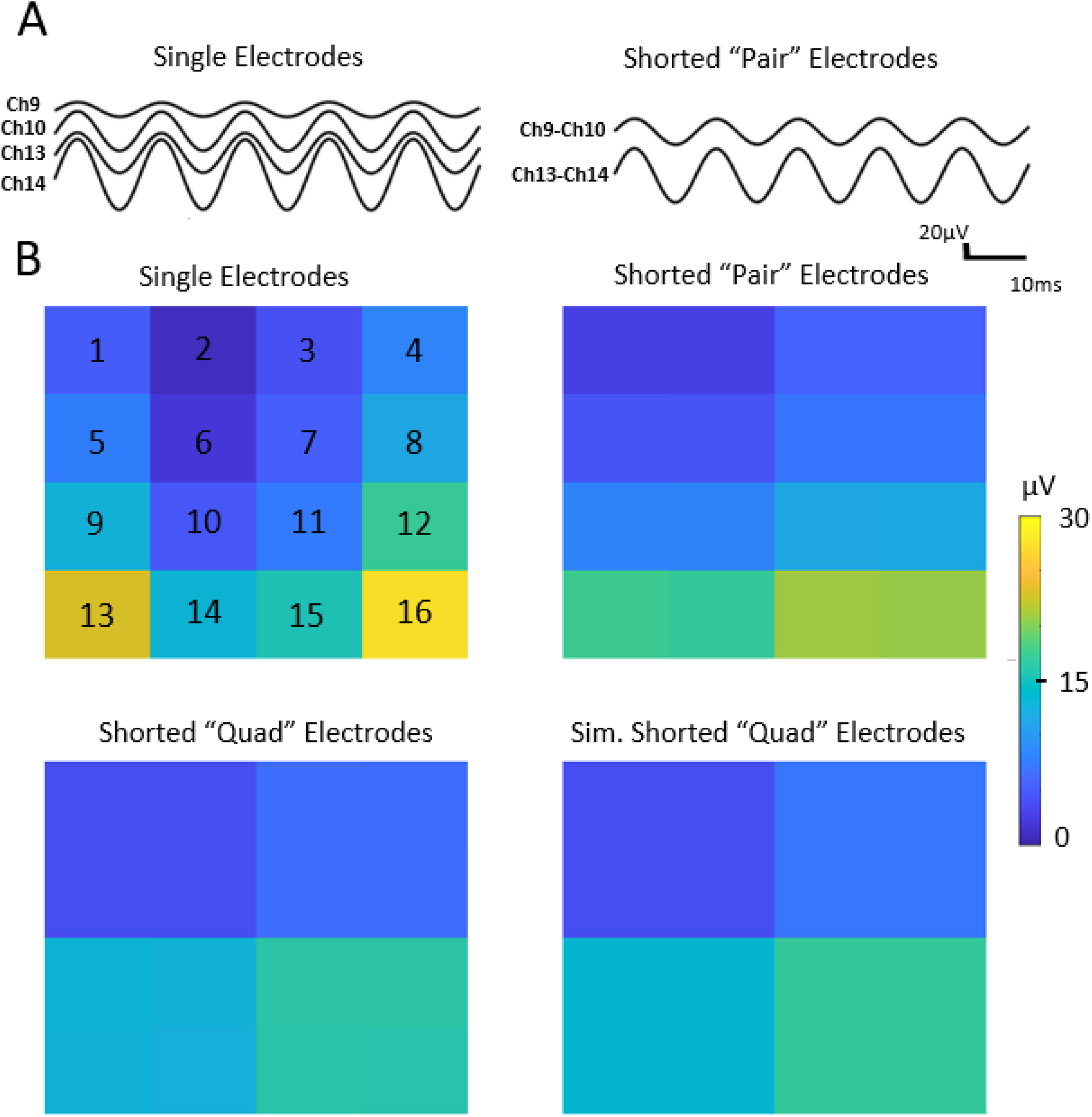
(A) Waveforms obtained for four single electrodes and the corresponding pair electrodes, e.g., Ch 9 and Ch 10 in the single configuration are shorted to obtain Ch9-Ch10 in the pair configuration. Similarly, Ch 13 and Ch 14 in the single configuration are shorted to obtain Ch13-Ch14. The single and pair recordings were obtained at different times, and the phases were aligned for the purpose of visual comparison. (B) Heat maps of RMS amplitude values measured using single, pair, and quad electrodes via physical shorting, as well as simulated quad electrodes via mathematical averaging (bottom right).

### 4.2 *In Vivo* Study

Data were collected from three human subjects (1 female, 2 male) aged 15.3, 6.1, and 19 years. Board-certified epilepsy specialists verified that the clinical interpretation of the ECoG study was unaffected by the multiscale recordings, and accurate localization of the seizure onset zone was achieved in all subjects (30).

#### 4.2.1 Correlation between channels is highest for large electrodes

In the human iEEG recordings, the correlation between channels at a fixed 6mm distance increased with electrode size in the low (1-30 Hz), gamma-1 (30-60 Hz), and gamma-2 (60-100 Hz) frequency bands in all three subjects (Figure 5 and Supplementary Figure 1). This increase was statistically significant when comparing small and large electrodes in the gamma-1 and gamma-2 bands in all three subjects (p<0.05, corrected for multiple comparisons). Moreover, the correlation values in all three frequency bands in all three subjects were significantly higher than the baseline correlation values (range: −0.02 to 0.01) calculated using the time shifted surrogate data.

**Figure 5:**
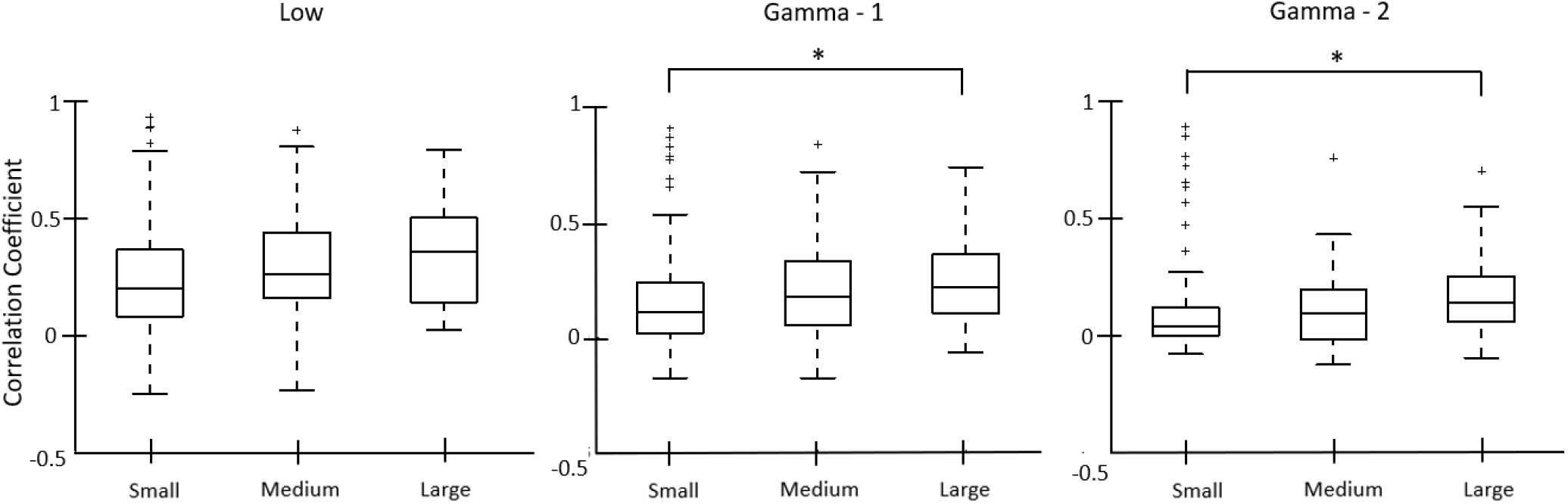
Boxplots of correlation values for electrodes at a fixed distance in each of the three electrode sizes (n=96 for small, n-=48 for medium, n=24 for large electrodes). The box plots are generated from data points corresponding to correlations in 100 five-second time windows from one subject. Results are shown for the low (left), Gamma-1 (middle), and Gamma-2 frequency bands (right). Data are shown for a single representative subject; data for the other two subjects can be found in Supplementary Figure 1. *indicates p-values< 0.05, ** indicates p<0.01, *** indicates p<0.001. All p-values are corrected for multiple comparisons using the Bonferroni method.

#### 4.2.2 Signal amplitude and power decrease with increasing electrode size

The iEEG RMS amplitude was highest for the small electrodes, and it decreased as electrode size increased (Figure 6A). The differences were statistically significant for all three electrode sizes across all three subjects.

**Figure 6:**
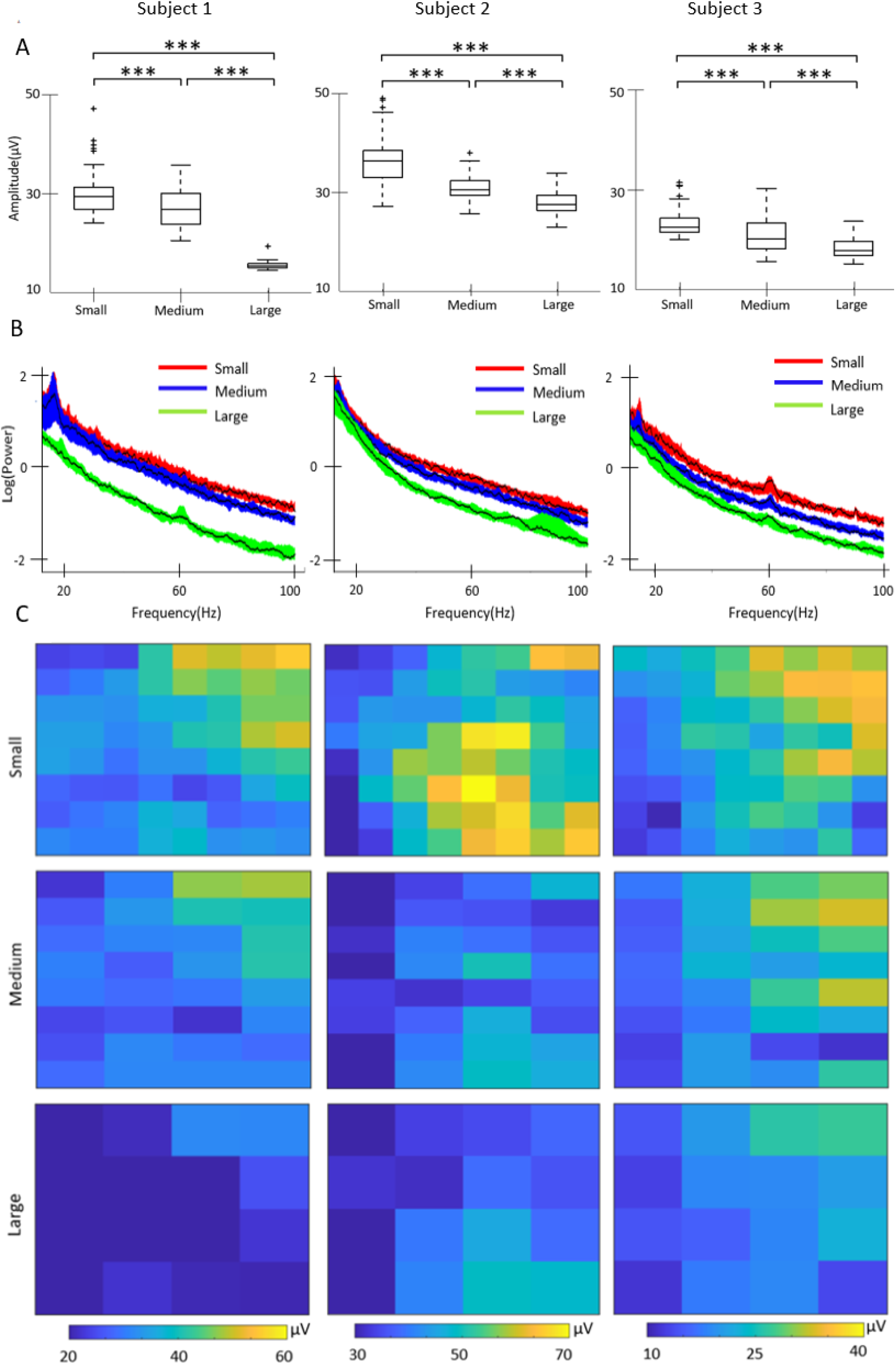
(A) Box plots of iEEG RMS amplitude in the 1-100 Hz frequency band measured using electrodes of three different sizes in the three subjects. Data from each subject is shown in a single column, across all subfigures. (B) Functional median curves for the power spectra in all three subjects. The black lines show the median curves, and the colored areas denote the 50% central region. (C) Heat maps of RMS amplitude values for all electrodes in each subject measured using small, medium, and large electrodes in the 1-100Hz band. For example, the results for the small electrodes show values in an 8×8 grid, with each colored square representing one electrode. Each column of subfigures corresponds to the results for one subject and the subfigures in each row correspond to small, medium and large electrodes. *indicates p-values< 0.05, ** indicates p<0.01, *** indicates p<0.001. All p-values are corrected for multiple comparisons using the Bonferroni method.

For the power spectra, the 50% central regions for the three electrode sizes overlapped in the low frequencies but were distinguishable for higher frequencies (Figure 6B), and the median periodogram curves were higher for small electrodes. This indicates that the iEEG power is higher for small electrodes compared to larger ones. Among the 48 sets of channels analyzed (16 from each subject), 39 showed a significant difference in the power spectra between the three different electrode sizes using the depth-based permutation test (p<0.05).

Note that there is significant spatial variation of the iEEG amplitude across the subdural grid (Figure 6C), likely related to differences between tissue inside and outside the seizure onset zone. However, for each patient, the spatial distribution of amplitudes was consistent across electrode sizes, with amplitude generally decreasing as electrode size increased. As in the *in vitro* experiment, the amplitude in the larger electrodes appeared to be consistent with the average of the corresponding small electrodes (Figure 6C).

#### 4.2.3 Interictal spike morphology depends on the size and location of the neural generator, relative to electrode size

To quantify the impact of electrode size on interictal spike morphology, we measured the spike SNR as a function of electrode size. We were unable to do direct event-wise comparisons using the physically shorted electrodes because the data for each electrode size were recorded at different times. Therefore, we used simulated spikes to estimate the change in SNR as electrode size varied. We first marked a total of 500 spikes, using the data from the small electrodes from all three subjects. For each spike, we then calculated simulated spikes in medium and large electrodes, via mathematical averaging of the associated small electrode iEEG.

Two examples of simulated spikes demonstrate why the smallest electrodes are not always associated with the highest SNR. In the first case (Figure 7A), the spike is clearly localized to a single electrode. Therefore, averaging reduces the SNR of the spike, and the biggest SNR is observed in the smallest electrode. In the second case (Figure 7B), the spike is more widespread, and averaging increases the amplitude relative to the background. Consequently, the largest SNR is observed for the medium electrode.

**Figure 7:**
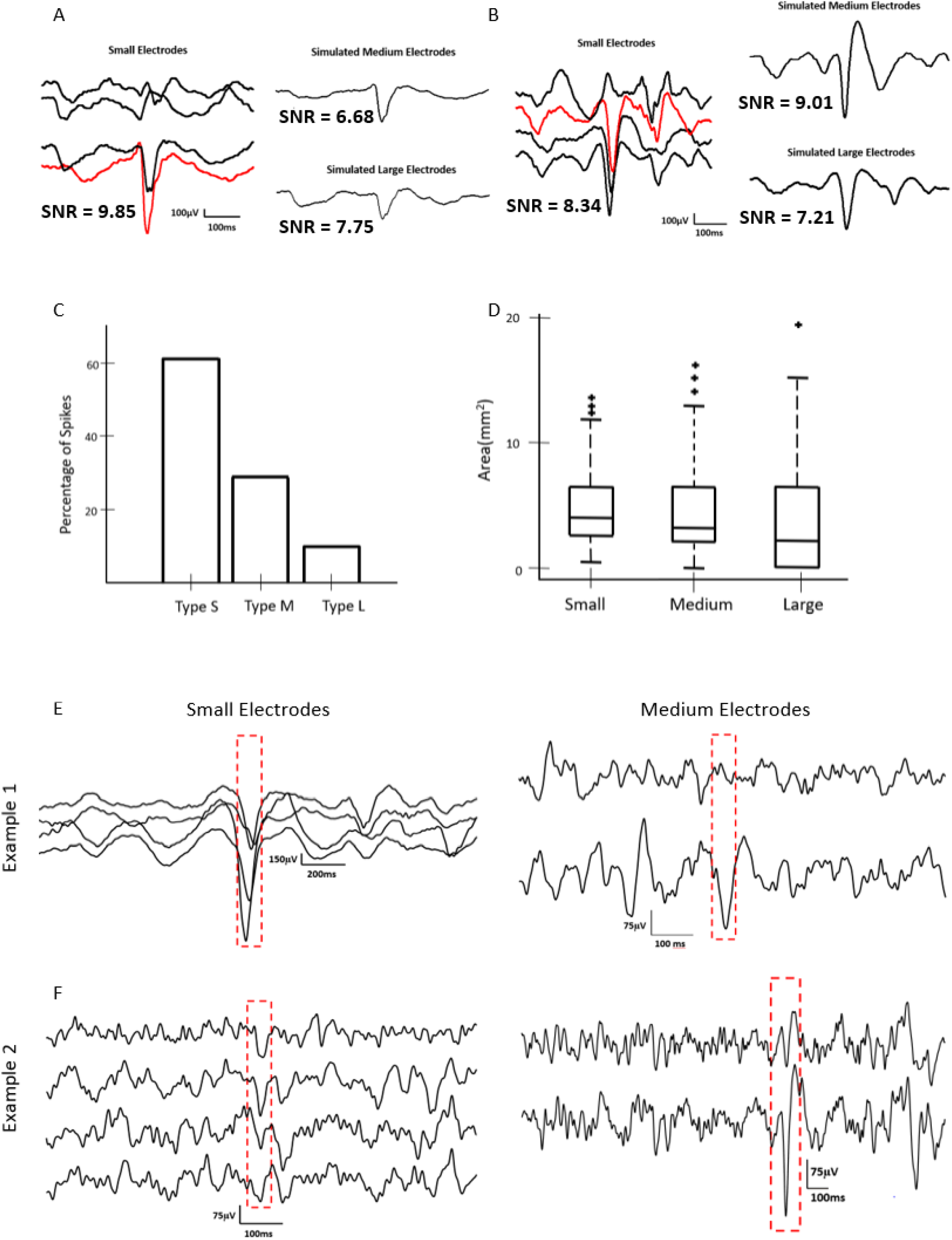
(A) Example of a type S interictal spike simulated via mathematical averaging, in which the SNR is highest for small electrodes. (B) Example of a Type M spike, in which the SNR is highest for simulated medium electrodes. (C) Bar graph showing the percentages of Type S, Type M, and Type L spikes based on simulated data for all three subjects. (D) Spatial spread of spikes for different electrode sizes across all three subjects. (E) Example of interictal spikes captured via physical shorting of electrodes, in which a high SNR spike is visible in the small electrodes (left) and a low SNR spike is seen in the medium electrodes (right). (F) A second example of an interictal spike captured via physical shorting of electrodes. Here, a spike with low SNR is visible in the small electrodes (left); when physical shorting was used to record from medium electrodes, larger SNR spikes were noted in the same set of channels (right).

Across all visually-marked spikes, 61% had the highest SNR in the small electrodes, 29% had the highest SNR in the medium electrodes, and 10% had the highest SNR in the large electrodes (Figure 7C). When measuring with small electrodes, the spikes were seen in a larger number of channels, but the cortical surface area of the spikes remained approximately constant across electrode sizes (Figure 7D, p>0.1).

Anecdotally, when visually comparing the interictal spikes from the three electrode sizes recorded via physical shorting, we found examples that were consistent with these results. We often found that the small electrodes were associated with the highest SNR (Figure 7E), but in some cases, the spikes were more visually prominent in the medium or large electrodes (Figure 7F).

## 5. Discussion

Through this study, we have introduced a method for dynamic selection of the size of iEEG electrodes after implantation in the human brain. We first presented an electrical circuit model, then performed an *in vitro* validation of that model, showing that there were no significant differences between physically shorted and mathematically averaged signals. In human subjects, we found that increasing electrode size leads to lower iEEG power and amplitude and higher correlation between channel pairs. The morphology of interictal spikes was also impacted by electrode size and depended on the size and location of the neural generator relative to the electrodes.

Our results support some hypotheses put forward in previous literature. In a study that measured correlation using both micro-ECoG and standard ECoG electrodes, the larger electrodes exhibited higher correlation (19). Wang et al. also reported higher degrees of dependence between larger electrodes, which is consistent with our results (Figure 5)(18). In an *in vivo* study of rat somatosensory cortex, the amplitudes of sensory evoked potentials (SEPs) recorded using small electrodes were higher than those recorded using larger electrodes (33). Our results in Figure 6 show the same trend. In our analysis of interictal spikes, we found that a majority of spikes had the highest SNR when recorded with small electrodes, but approximately 40% of spikes had the highest SNR when recorded with medium or large electrodes. Anderson et al. (2010) posited that using larger electrodes for recording action potentials of neurons decreases the SNR. Assuming that the spatial extent of the action potential is small relative to the electrode size, this is consistent with our observations of interictal spikes (Figure 7)(15). While single neuron spikes occur on a much smaller spatial scale than interictal epileptiform discharges, the two cases share conceptual similarities.

There are some limitations to this study. The electrodes in the ECoG grids had a center-to-center distance of 3mm, so the adjacent shorted electrodes were not contiguous. That is, there was some area of tissue between shorted electrodes that was not in contact with them, which is a deviation from the assumption of a single, continuous electrode. Using tripolar concentric ring electrodes (34) or decreasing the interelectrode distance in the grid would help alleviate this limitation. However, because this study was done on human subjects, we were limited to the use of FDA approved electrodes. Another limitation was that the recordings from electrodes of different sizes were done sequentially and, therefore, occurred at different times. We repeated these measurements to verify that our findings remained robust and were stable over time. In particular, we found no significant difference in the value of the power spectrum when comparing data spaced 10 minutes apart, for any electrode size. Additionally, the data in this study were obtained from patients with epilepsy, and this disease is known to alter various features of the iEEG data. Because our aim was to study the effects of electrode size on iEEG, irrespective of the origin of the activity, we believe this did not significantly impact our results. Moreover, all comparisons were made using data recorded from the same region of brain tissue; therefore, the presence or absence of epileptogenic activity should impact all conditions equally.

Future work in this field may benefit from the use of modeling techniques like Finite Element Modeling (FEM). FEM based methods have been used to numerically solve the EEG forward problem accurately, that is, to determine the voltages at the surface of the brain given the location of deep sources. This is done by incorporating complex geometries and electrical properties of the brain into the model (35). FEM could be applied to our study to exactly simulate electrodes of different sizes having similar geometries and spacing between them. This would address the inconsistency between our simple mathematical averaging approach and the non-contiguous area of the larger electrodes in this study. Thus, an FEM-based approach could provide a more accurate mathematical model of the measured electrical activity, as a function of electrode geometry and spacing, given a set of neural sources.

This study is the first to present a method to record intracranial EEG from a static section of neural tissue using electrodes of different effective sizes. This technique provides an avenue for multi-scale analysis of neurological phenomena recorded from a single location in the brain. The methods used here could also enable dynamic selection of optimal electrode sizes for detection of neurological events like seizures, HFOs, and interictal spikes, as well as recordings used by neural prostheses or BCIs. This is especially relevant in a clinical setting where the precise locations of these events are unknown prior to surgery and are highly variable across patients. Clinicians rely on visual analysis of the iEEG, and electrographic events that are barely visible in data from a particular electrode size could be more accurately studied when measured with a larger or smaller electrode. In applications like neuroprostheses and BCI where the quality of the signals is paramount, our methods can be used to maximize SNR while requiring only a single implantation of a standard commercially available electrode grid. Overall, this technique has the potential to facilitate patient-specific optimization of iEEG recordings, for both clinical and engineering applications.

## Acknowledgements

Research reported in this publication was supported by the National Institute of Neurological Disorders and Stroke of the National Institutes of Health under award number R01NS116273 and a CHOC PSF Tithe grant. The content is solely the responsibility of the authors and does not necessarily represent the official views of the National Institutes of Health. We would also like to acknowledge Zoran Nenadic and Jeffrey Lim for their support and helpful feedback, particularly in the early stages of this work.

