## Appendix for "A novel method for multi-scale recording of intracranial EEG through dynamic alteration of electrode surface area"

### Circuit Model Calculations

Please refer to Figure 1B for a diagram of the electrical circuit.

The voltage observed at the brain surface ($V_{b1}$ and $V_{b2}$) due to two sources $V_{s1}$ and $V_{s2}$ can be calculated from the circuit, considering only the part to the left of the red line. We get the voltages to be:

$$V_{b1} = \frac{Z_{b2} + Z_{12}}{Z_{b1} + Z_{b2} + Z_{12}} V_{s1} + \frac{Z_{b1}}{Z_{b1} + Z_{b2} + Z_{12}} V_{s2}$$

$$V_{b2} = \frac{Z_{b2}}{Z_{b1} + Z_{b2} + Z_{12}} V_{s1} + \frac{Z_{12} + Z_{b1}}{Z_{b1} + Z_{b2} + Z_{12}} V_{s2}$$

When the recording electrodes are added to the circuit (right of the red line), the voltages sensed at each electrode ($V_{e1}$ and $V_{e2}$) will be approximately equal to $V_{b1}$ and $V_{b2}$ respectively, considering the input impedance of the amplifier to be very large.

Then, for example, if we assume equal tissue impedances $Z_{b1}, Z_{b2}$ and $Z_{12}$, the voltage sensed at each electrode would be,

$$V_{e1} = \frac{2}{3} V_{s1} + \frac{1}{3} V_{s2}$$

$$V_{e2} = \frac{1}{3} V_{s1} + \frac{2}{3} V_{s2}$$

To represent an electrode with twice the surface area, we model two adjacent electrodes that are shorted together (green dashed line). In this case, the voltages at the brain surface are

$$V_{b1} = \frac{Z_{b2}(Z_e + Z_{12}) + Z_{12}Z_e}{(Z_e + Z_{12})(Z_{b1} + Z_{b2}) + Z_{12}Z_e} V_{s1} + \frac{Z_{b1}(Z_e + Z_{12})}{(Z_e + Z_{12})(Z_{b1} + Z_{b2}) + Z_{12}Z_e} V_{s2}$$

$$V_{b2} = \frac{Z_{b2}(Z_e + Z_{12})}{(Z_e + Z_{12})(Z_{b1} + Z_{b2}) + Z_{12}Z_e} V_{s1} + \frac{Z_{b1}(Z_e + Z_{12}) + Z_{12}Z_e}{(Z_e + Z_{12})(Z_{b1} + Z_{b2}) + Z_{12}Z_e} V_{s2}$$

Where $Z_e$=$Z_{e1}$+$Z_{e2}$.

Note that the expressions for $V_{b1}$ and $V_{b2}$ become approximately equal to the un-shorted case when the electrode impedance is small compared to the tissue impedance.

Finally, the common voltage sensed by the two shorted electrodes ($V_{e1}$=$V_{e2}$=$V_e$) is given by:

$$V_e = \frac{Z_{e2}}{Z_{e1} + Z_{e2}} V_{b1} + \frac{Z_{e1}}{Z_{e1} + Z_{e2}} V_{b2}$$

If we assume equal electrode impedances $Z_{e1}$= $Z_{e2}$, then we obtain,

$$V_e = \frac{V_{b1} + V_{b2}}{2}$$

This is the average of the two voltages from the individual electrodes before shorting. Therefore, this model suggests that physical shorting of adjacent electrodes is equivalent to mathematical averaging of the activity recorded by the individual electrodes.
