## Supplementary Figure 1 for "A novel method for multi-scale recording of intracranial EEG through dynamic alteration of electrode surface area"

### Supplementary Figures

#### Correlation between channels

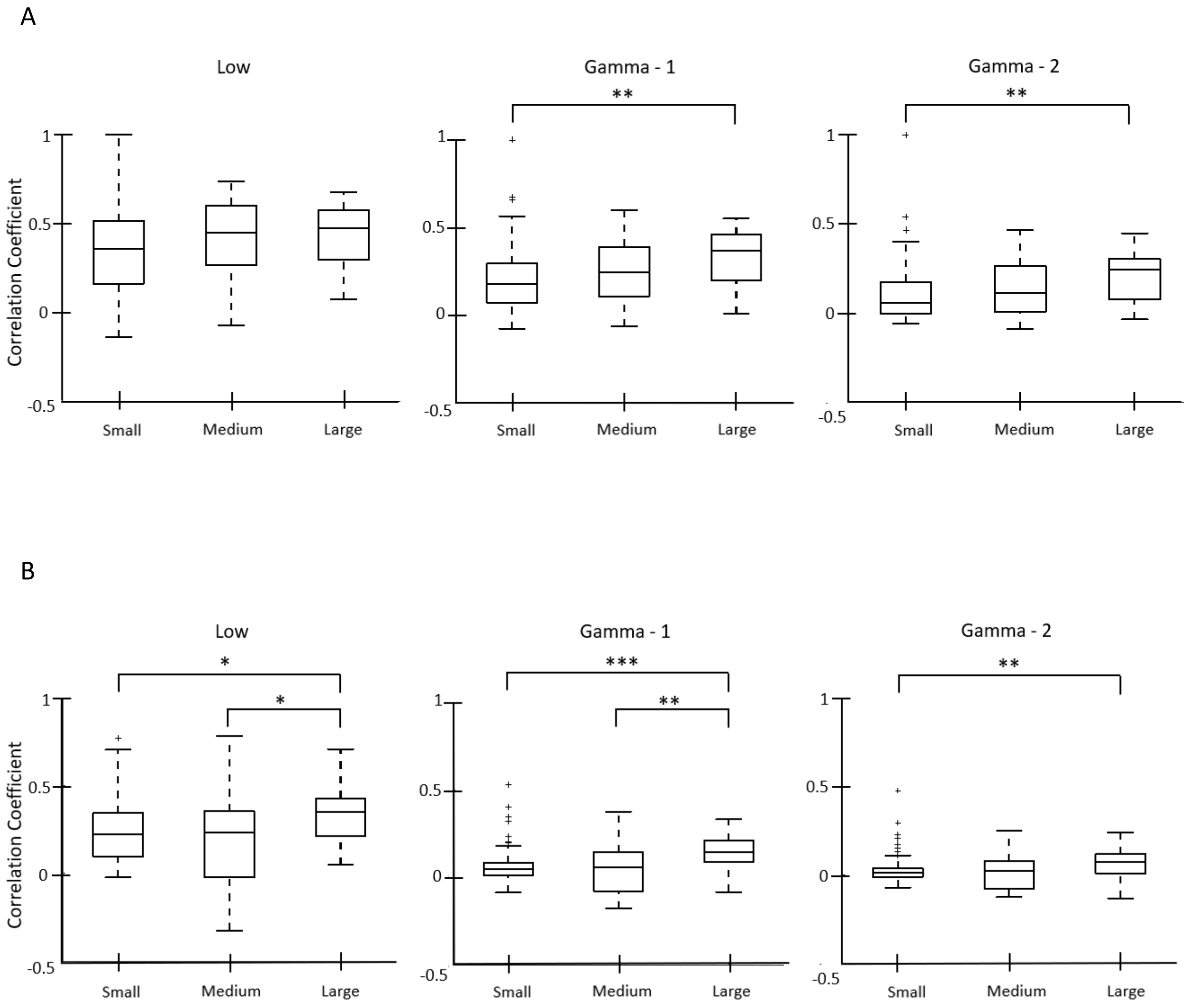


*Supplementary Figure 1: Boxplots of correlation values for electrodes at a fixed distance in each of the three electrode sizes for the three frequency bands – low, gamma 1 and gamma 2. A and B show the other two subjects from the study, as a complement to Figure 5.*
